# A multi-session simultaneous EEG-fMRI dataset with repeated experience sampling

**DOI:** 10.64898/2026.02.04.703882

**Authors:** Lotus Shareef-Trudeau, David Braun, Tiara Bounyarith, Janet Z. Li, Huiling Peng, Aaron Kucyi

**Affiliations:** Department of Psychological and Brain Sciences, Drexel University, Philadelphia, PA, USA; Temple University Brain Research & Imaging Center, Temple University, Philadelphia, PA, USA

## Abstract

The integration of electroencephalography (EEG) and functional Magnetic Resonance Imaging (fMRI) can be used to characterize temporal and spatial components of neural activity during unfolding mental experience. Here we introduce a multi-session simultaneous EEG-fMRI dataset with measures of continuous behavior and spontaneous mental experience. Data components, organized in Brain Imaging Dataset Structure (BIDS) format, include fMRI, EEG with carbon wire loop sensors for artifact removal, continuous performance task responses, experience sampling ratings, and mental health surveys, from 24 healthy adults. Tasks included the gradual-onset continuous performance task and resting state with intermittent experience sampling of 13 unique thought dimensions (36 repetitions, including 468 total ratings, per participant). The same protocol was completed on two different days, yielding approximately 1.33 hours of simultaneous EEG-fMRI data per individual. The dataset may be used to explore the behavioral and experiential relevance of brain activity during the wakeful resting state. The dataset also provides a means to study the reliability of relationships between fMRI and EEG features across sessions within individuals.

## Background & Summary

Researchers have increasingly become interested in studying the neural bases of spontaneous, stimulus-independent, and off-task thought (often collectively considered as the study of “mind-wandering”)^1–3^. Off-task thought has been estimated to occur during nearly half of a typical individual’s waking hours^1,2^ and has been linked to mental functions including cognitive control, creativity, memory, and mental health^3–7^. Here we introduce a simultaneous electroencephalography (EEG) and functional magnetic resonance imaging (fMRI) dataset^8^ to provide the opportunity to investigate the neural correlates of mind-wandering, attentional fluctuations, and spontaneous mental experience with high spatial and temporal resolution. Few studies have employed simultaneous EEG-fMRI methodologies to investigate mind-wandering^9^, and, in contrast to our study, none to our knowledge have done so employing a resting state condition with repeated experience sampling.

Many researchers have used EEG-fMRI to study neural activity during resting states—or conditions where participants are not given an explicit task other than to stay awake. These methodologies have provided insight into the electrophysiological basis of spontaneous activity in functional networks that are commonly identified in fMRI data^10–13^. It is notable, however, that at rest, people engage various spontaneous cognitive, emotional, and perceptual processes^14,15^. As such, resting state paradigms may provide privileged, naturalistic information about brain functional organization across the various mental states that are active when attention is not focused on a task. However, for neural insights to be linked to observable behavior, it is necessary to collect responses on a behavioral task^16^. Repeated experience sampling, conducted online during neural recording, is one commonly used approach for mapping specific momentary conscious experiences to corresponding brain states in EEG^17,18^ and fMRI^19^ studies. Multidimensional experience sampling (MDES) is specifically useful for measuring and modeling how different contents and processes of thought group and/or change together^20,21^.

Past studies have employed repeated experience sampling online during resting state fMRI^22–24^, retrospective experience sampling after resting state fMRI^25–27^ and repeated experience sampling online during rest^28^ and naturalistic tasks^29^ during EEG. One recent EEG-fMRI study investigated mind-wandering (defined as task-unrelated thought) during sustained attention tasks using unidimensional experience sampling^9^. Relative to these past works, our approach of combining repeated MDES at rest during simultaneous EEG-fMRI presents a novel approach to investigating spontaneous brain activity and mental dynamics.

In our MDES paradigm, participants each provided multiple ratings per trial to describe their thoughts during rest across two EEG-fMRI sessions. Using these reports, we aimed to provide a close approximation of individuals’ subjective experience whilst maintaining standardized reports across trials and participants^30,31^. In addition to resting state MDES (rs-MDES), participants completed the gradual-onset continuous performance task (GradCPT), a paradigm where participants make go/no-go responses to gradually morphing stimuli^32^. This combination of rs-MDES and GradCPT allows investigation into how behavioral and subjective measures of attention may carry independent and/or interrelated information across conditions.

An important feature of our dataset is the use of carbon wire loop (CWL) sensors in our EEG caps during recordings. The simultaneous EEG-fMRI method has developed substantially over the past 30 years^33^, including technological developments to optimize participant safety and data quality^34,35^. However, combining these two modalities presents methodological challenges. Recording EEG during fMRI acquisition introduces major artifacts in the EEG data, primarily MR gradient and cardioballistic artifacts, that cannot be fully corrected with typical EEG preprocessing techniques. Moreover, EEG artifacts can arise from the scanner’s helium pump which produces vibrations in the scanner bore^36,37^. Recent hardware innovations, such as CWLs, are capable of measuring these systematic artifacts, which enables regressing out these artifacts during preprocessing^37,38^.

## Methods

### Participants

Simultaneous EEG-fMRI data were collected in 24 adults (ages 18-35 years; mean age: 21.17 years, 64% female) recruited from the Philadelphia, Pennsylvania area, primarily drawing from communities in and around Drexel University and Temple University. Participants were fluent in English, right-handed, and had normal/corrected-to-normal vision. Potential participants were excluded from the study if they were unable to provide consent, reported having a current or history of psychiatric/neurological disorders, a chronic medical condition, were pregnant, were prisoners, were unable to understand English, had metal in their body, were contraindicated for MRI, had allergies to saline gel, or had cold, flu or COVID-19 symptoms within the two weeks preceding participation. Two participants were excluded because they were unable to fit into one of our three EEG cap sizes (54cm, 56cm, and 58cm circumference). We used a 64-channel head coil for MRI, a relatively small head coil, that could not accommodate MR-safe glasses. Participants who wore glasses and did not have access to contact lenses were screened out during recruitment. One of our participants completed only one of the two scheduled EEG-fMRI sessions due to scheduling conflicts. Our total dataset comprises 47 EEG-fMRI sessions (2 sessions each for 23 subjects, 1 session for 1 subject).^8^

### Overview of experimental procedures

Participants attended three sessions at the Temple University Brain Research & Imaging Center with time between the two scanning sessions (i.e., the last 2 visits) ranging from 3 to 62 days (mean = 12.61 days, median = 7 days). The study protocol was approved by the Drexel University Institutional Review Board (IRB Protocol #: 2208009389). All methods were carried out in accordance with approved protocol.

In a first session prior to the main experiment, written informed consent was obtained from each participant alongside demographic information. Participants then filled out a series of questionnaires pertaining to different mental health conditions, including a crosscutting assessment: DSM-5 Self-Rated Level 1 Cross-Cutting Symptom Measure-Adult (DSM XC)^39^, a measure of impairment: WHO Disability Assessment Schedule-12 (WHODAS-12)^40^, a measure of anxiety: General Anxiety Disorder-7 (GAD-7)^41^, measures of depression: Patient Health Questionnaire-9 (PHQ-9)^42^, Ruminative Response Scale (RRS)^43^, and a measure of trait mind-wandering: Mind-Wandering Deliberate-Spontaneous scale (MWD-S)^44^. All questionnaires were combined and administered online through REDCap electronic data capture tools^45,46^. During the first session, participants were trained on all experimental tasks and informed of the full EEG-fMRI study procedure for upcoming scanning days (sessions two and three). Additionally, head circumference was measured from glabella to inion to determine EEG cap size to ensure that we had the proper fit for them. The first session lasted one hour.

On scanning days, measures of state anxiety were obtained using the state scales of the State Trait Anxiety Inventory (STAI-S^47^, and participants provided an estimation of number of hours slept the night before. Scanning sessions on visits 2 and 3 ran for a total of 2 hours each (see **Figure 1a**), with approximately 1-hour dedicated to setup of the EEG system and MRI safety procedures. Prior to cap setup, subjects were checked for metal using first a metal detector wand and then using a Ferroguard standing detector. Once confirmed that all metal items had been removed, participants were seated. Alcohol wipes were used to clean the forehead of oil and debris, and then the appropriate EEG cap was placed on the head with Fp1 and Fp2 electrodes aligned directly above the line of the eyebrows and secured with washer stickers. The cap was tugged and smoothed to remove all bubbling, and a secure Velcro strap was attached under the chin of each participant. All electrodes were then filled with electrolyte gel.

**Figure 1.**
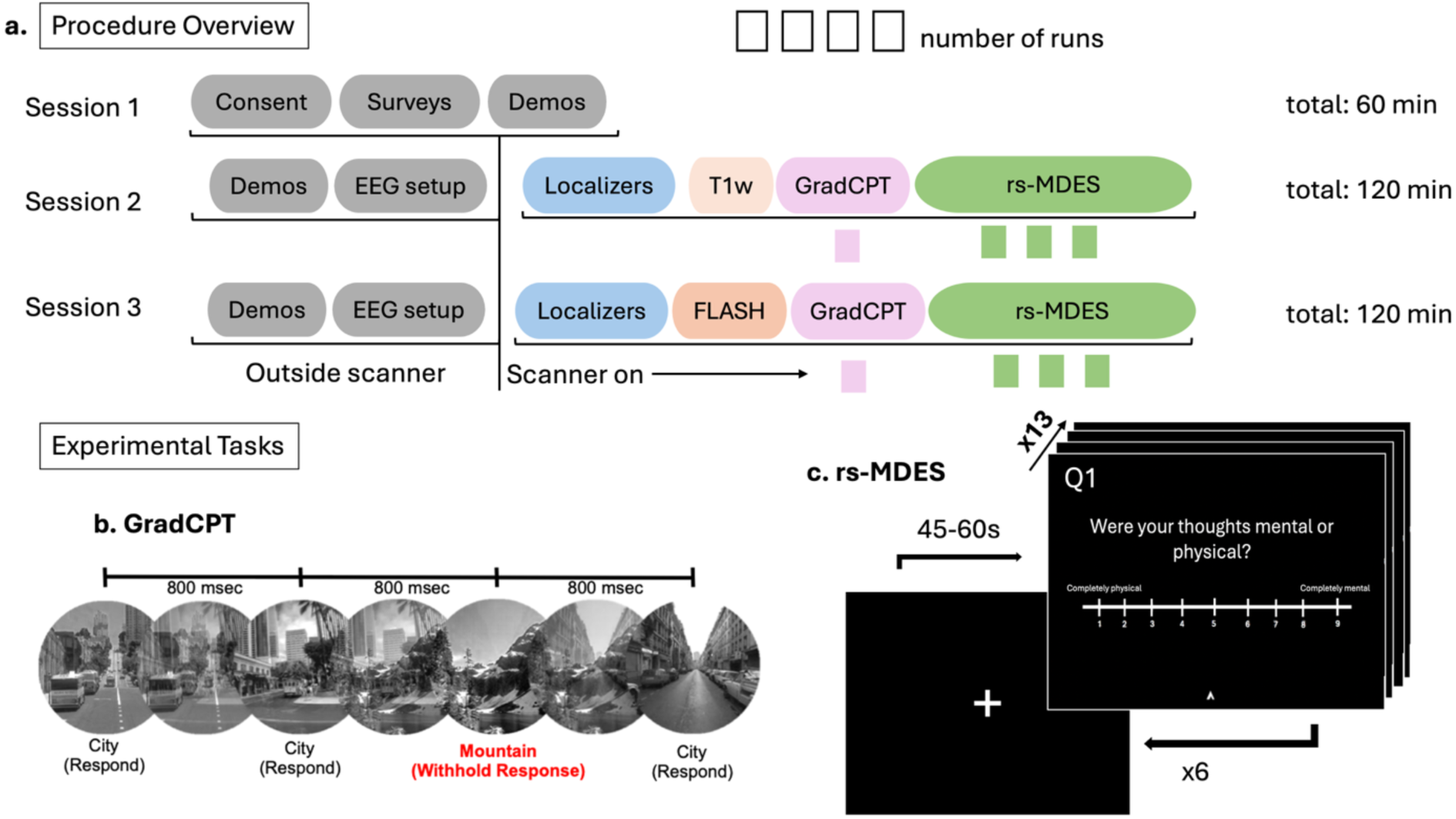
Overview of study procedures and tasks. **a)** Study procedure for sessions 1-3. Tasks that took place outside of the scanner are indicated in grey. Session 1 was a purely behavioral orientation session that ran for 1 hour. Sessions 2-3 (scanning days), lasted two hours in total with half of the session dedicated to out-of-scanner procedures and the other half to simultaneous EEG-fMRI collection. In the first hour, participants underwent task training, MRI safety preparations, and EEG set up. The second hour was dedicated to simultaneous EEG-fMRI data collection. The scanning protocol included collection of localizers and a T1 weighted Magnetization Prepared Rapid Gradient Echo (MPRAGE) structural scan. Functional MRI scans were collected across two tasks, GradCPT and resting state multidimensional experience sampling (rs-MDES). Session 3 was identical to Session 2 excluding the structural scan, which was Fast Low Angle Shot (FLASH) instead of MPRAGE sequence in Session 3. **Tasks. b)** The GradCPT was run for 8 minutes during which participants were instructed to respond with a button press to frequent city scenes and withhold response to the presentation of rare mountain scenes. **c)** Three runs of the rs-MDES were collected at each session wherein participants visually fixated on a cross at the center of their screen and let their minds-wander. Intermittent thought probes appeared pseudorandomly every 45-60 seconds which were rated by participants at their own pace. Thirteen probes were presented in sequence, capturing unique dimensions of subjective experience for 6 trials across the duration of each run (3 rs-MDES runs per session). At the end of each run, the scan was manually stopped, producing data files which vary in total duration.

### EEG acquisition

EEG data were collected using a MR-compatible system (Brain Products Gmbh, Gilching, Germany), including two amplifiers (BrainAmp MRPlus and BrainAmp ExG) and the BrainCap MR with four CWL sensors embedded. Additional EEG-fMRI specific equipment included a Brain Products PowerPack (MR-compatible battery pack for the amplifiers), Triggerbox (sends and receives MRI and stimulus triggers to EEG recording software), a SyncBox interface and main unit (synchronizes EEG acquisition phase with MR clock), a BrainVision USB 2 Adapter (connection hub for all EEG components), and an MR sled (mobile surface for mounting the EEG components into the scanner bore). Recorded channels included 4 CWLs, 31 cortical channels and one electrocardiography (ECG) channel (channel 32) placed on the back. In addition, the cap also contained a reference and ground electrode at locations FCz and AFz respectively. The cap’s built-in serial current-limiting resistors included 15kOhms of resistance for the ground and reference electrodes, 10kOhms of resistance for all 31 EEG electrodes, and 20kOhms of resistance for the ECG electrode. All electrodes were connected to the BrainAmp MRPlus. The four CWLs sit over the frontal left, frontal right, parietal left, and parietal right locations on the cap and were connected to the bipolar BrainAmp ExG. The CWL channels record minute head movements in the scanner environment, used to perform advanced cardioballistic, helium pump, and movement artifact correction^38^ (see **Figure 2a and 2b**). Cortical electrodes were arranged according to the international 10-20 system. Electrodes were filled using high-chloride abrasive electrolyte-gel (Abralyt HiCl). A layer of Surgilast, an elastic dressing retainer, was placed over the cap after set-up to ensure electrode contact with the scalp and minimize cap movement. Electrode impedance was kept below 20kOhms and was verified twice, once during set up, then inside the scanner just before the closing of the head coil. The EEG data were recorded using BrainVision Recorder V.1.25.0201 software at a sampling rate of 5kHz on a Windows operating system using a Dell Latitude 7430 64-bit operating system with a 12^th^ Gen Intel Core i7-1265U and 16GB of Ram.

**Figure 2.**
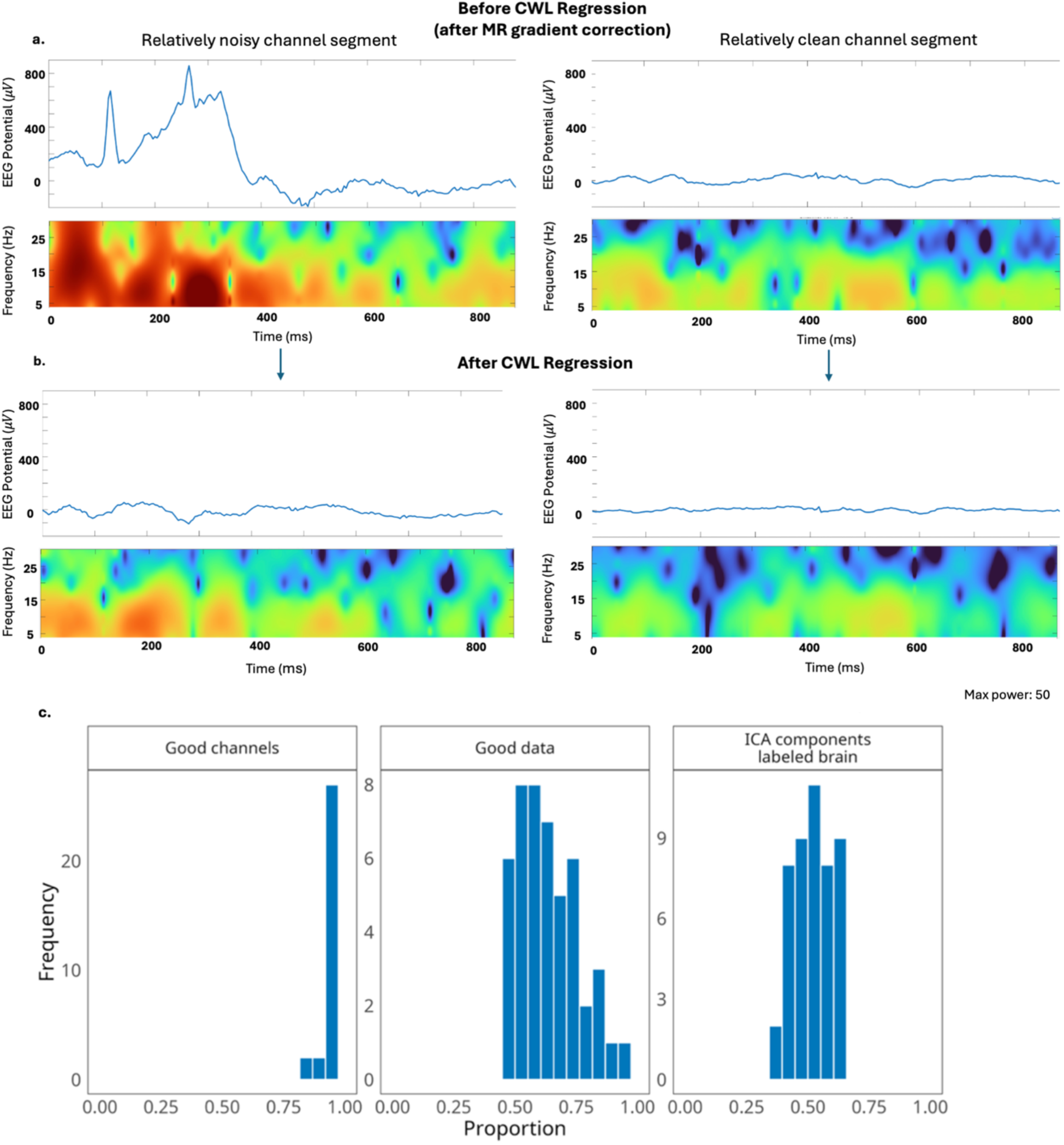
EEG artifact correction and quality check. **a)** Example of two 900-ms windows of CWL-based correction of EEG data recorded during simultaneous EEG-fMRI. On the left, a relatively noisy channel channel segment is shown (sub-009 session-001 run-001) and on the right, a relatively clean channel segment is shown (sub-016 session-002 run-001). Both examples appear on the same scale for the y (-200 to 900 microvolts) and x (0-900ms) axes and at the same electrode, P8. **b)** Example of the same 900-ms windows of EEG data after CWL signal regression. **c)** Figures show distributions of percent ‘good channels’, percent of ‘good data’ and percent of independent components *identified as sources of brain activity. These analyses and figures were based on Delorme et al.*^48^ *(see main text for descriptive details)*.

### Simultaneous EEG-fMRI recording setup

After cap set up, the EEG-fMRI equipment was set up in the control and scanning rooms. The TriggerBox was attached to the fMRI button response system for communication between the EEG equipment, our stimulus software, and the fMRI triggers. Using the TriggerBox, we set a 5ms 15 bit stretch of the fMRI pulse to allow for our system to process the trigger. The fMRI volume markers, labeled as “T 1” events, were therefore recorded in the EEG data at a 5ms offset from when they were sent. The Syncbox was routed from the equipment room to the control room and delivered timing information from the MR clock to the EEG system, maintaining phase synchrony. Sync status markers were recorded in the EEG data every two seconds, where “SyncStatus/Sync On” event labels indicated successful synchronization between the MR and EEG systems. The Powerpack, ExG amp and BrainAmp MR were placed in the bore of the scanner in a vertical stack on an MR sled. Then the amplifiers were connected to the USB 2 adapter (the connection hub for all components of the EEG system) in the control room via fiberoptic cables. Participants were carefully positioned on the scanner bed to minimize electrode disturbance. Cushions and airbags were placed around the head to minimize head motion during the scan. A belt was looped around the MR sled and beneath the head coil to attach the two for stability. The EEG ribbon cables were routed from the top of the head, through a channel in the base of the head coil and attached to the amplifiers in the scanner bore. A sandbag was placed on top of the cables to minimize vibrations.

For task response, participants held a MR-safe Current Designs 4-finger button box in the right hand. A screen displaying visual stimuli was rear projected using a Psychology Software Tools Hyperion projector at the end of the scanner bore. Separate EEG data recordings were taken for each run of each fMRI task with a variable delay between onset of EEG recording and initiation of fMRI acquisition. The EEG data received event markers from the stimulus computer and volume triggers for each acquired fMRI frame from the Siemens MRI console. Each event triggered two successive markers automatically named by the TriggerBox. The first event marker appears inconsistently named across participants, while the second marker name stayed consistent across tasks and participants, named “Stimulus/S255.”

### MRI data acquisition

MRI data were acquired using a 64-channel head coil on 3.0 Tesla Siemens MAGNETOM Prisma. To accommodate the EEG-fMRI set-up, the 64-channel head coil included an aperture for routing cables from the top of the EEG cap into the scanner bore to connect to EEG amplifiers. The first 22 subjects and the first session of the 23^rd^ subject were run on Siemens software version Syngo MR VE11E. The second session of the 23^rd^ subject and both sessions of the 24^th^ subject were run on software version Syngo MR XA30. All acquisition parameters remained consistent across both software versions with the exception of nonlinear gradient correction, set to “false” for the first 22 subjects and set to “true” after the software upgrade. This change reflected differences in the default settings of the scanner rather than a deliberate change made by the researchers. After localizer scans, a 3D high resolution MPRAGE structural T1-weighted image was acquired (TR = 2400 ms; TI = 1000 ms; TE = 2.22 ms; slices = 208; FoV = 256 x 240mm, voxel size = 0.8 mm^3^ isotropic; flip angle = 8°; partial Fourier off; pixel bandwidth = 220Hz/Px) during the first scanning session. During a participant’s second scanning session, a fast T1-weighted, 2D MRI sequence was collected (TR = 190 ms; TE = 2.46 ms; in-plane voxel size = 0.8 x 0.8mm, slices = 25; slice thickness = 4.00 mm; flip angle = 70°; pixel bandwidth = 280 Hz/Px). Following the T1-weighted scan, we ran a B0 field map for the purpose of correcting echo-planar imagining (EPI) fMRI images (TR = 789 ms; TE 1 = 4.92 ms; TE 2 = 7.38 ms; slices = 81; FoV = 192mm x 192mm; voxel size = 2.0 mm^3^ isotropic; flip angle = 45°; partial Fourier off; pixel bandwidth = 668 Hz/Px). All T2* (BOLD fMRI) sequences were acquired with a CMRR multiband gradient-echo EPI single shot sequence aligned with the AC-PC plane (TR = 2000 ms; TE = 25.00 ms; flip angle = 70°; slices = 81; FoV = 192mm x 192mm; voxel size = 2.0 mm^3^ isotropic; multi-band acceleration factor = 3; phase encoding direction = anterior-to posterior). A total of four fMRI runs across two tasks (one GradCPT run, three rs-ES runs) were run for 46 out of 47 scanning sessions, where one session (sub-003_ses-001) was ended early due to time constraints and only 3 runs were collected.

### Experimental Tasks

In this section, we describe the task paradigms and behavioral data collected during simultaneous EEG-fMRI. All tasks were designed and administered in Psychophysics toolbox version 3.0.18^49^ in MATLAB R2023b and run on an Alienware laptop running Windows 11 operating system.

*Gradual Onset Continuous Performance Task* (GradCPT; 1 run per session, 8 minutes). The GradCPT was designed to engage sustained attention and track attentional fluctuations^32^. During this task, participants viewed a stream of grayscale images (cities, mountains) and were instructed to respond with index finger button presses to city scenes while withholding responses to mountain scenes (**Figure 1b**). The images were presented in a randomized order and quickly transitioned from one image to another approximately every 800 ms in a continuous stream for 600 trials (via linear pixel-by-pixel interpolation between consecutive images). The frequent city scene stimuli were programed to be presented on ∼90% of trials and mountain scene stimuli were presented on ∼10% of trials. Data on accuracy of button presses in response to stimuli and precise reaction time information were recorded using an algorithm which determines whether a button press belongs to the previous or current trial (for a more detailed description of the algorithm rules, see Esterman et al.)^32^.

*Resting State Multidimensional Experience Sampling* (rs-MDES; 3 runs per session, ∼10.5 minutes each). We designed the rs-MDES task to naturalistically study the content and dynamics of ongoing subjective experience^28^. Participants completed three runs per session, each comprising six trials of a resting state period followed by MDES. During each resting state period, participants viewed a central fixation cross and were instructed to let their minds wander (**Figure 1c**). After a random integer time interval between 45-60 seconds following each onset of the fixation cross, 13 distinct probes appeared in sequence. Each probe was presented with a Likert scale ranging from 1-9 including anchors describing the extremities. Participants could advance through the questions at their own pace, provided they responded within the 12 second time limit that was placed on each question. If a participant did not respond within the time limit for any given question, the task would automatically move on to the next probe and record a failed response. Responses to probes were made using the MR button box, where the index finger button was used to move the arrow on the scale to the left, the middle finger button was used to move the arrow to the right, and the ring finger button was used to enter the response and advance to the next question. The pinky finger button was used as an extra option for the first question under special circumstances described below. Participants were instructed to answer as quickly and accurately as possible. During session one, participants were trained on definitions and guided through several example trials before the task began. Because this task was largely self-paced the duration of each run varied such that 6 trials, with each including 13 ratings, were always completed per run. Total run time ranged from 7 minutes 46 seconds to 13 minutes 41 seconds (mean: 10 minutes 28 seconds; SD: 1 minute 10 seconds). To accommodate the self-paced nature of this task, the scanner was stopped manually at the end of each run, resulting in a different number of fMRI volumes (and EEG recording durations) from run to run. Across all sessions and all participants, experience sampling data from 840 trials were collected. Our approach to experience sampling was similar to that employed in other mind-wandering studies^7,28,50–52^. The 13 probe questions and scales that appeared are listed below:

1. Stimulus-independent attention: “Were you more focused on your own thoughts (mental) or on sensing something in the world or in your body (physical) or was your mind blank?” (1=completely physical, 9=completely mental, or pinkie button press to indicate mind blank)
2. Temporality-Past: “Were your thoughts oriented towards the past?” (1=not about the past at all, 9=completely past oriented)
3. Temporality-Future: “Were your thoughts oriented towards the future?” (1=not about the future at all, 9=completely future oriented)
4. Self-Referential: “Were your thoughts about yourself?” (1=nothing about you, 9=completely about you)
5. Regarding others: “Were your thoughts about others?” (1=nothing about others, 9=completely about others)
6. Arousal: “How activated or energized were you feeling?” (1=completely deactivated, 9=completely activated).
7. Affect: “How positive or negative did you feel?” (1=completely negative, 9=completely positive)
8. Engagement: “How easy was it to disengage from your thoughts?” (1=extremely easy, 9=extremely difficult)
9. Intentionality: “How intentional were your thoughts?” (1=completely unintentional, 9=completely intentional)
10. Movement: “Were your thoughts freely moving?” (1=unmoving, 9=moving freely)
11. Imagery: “Were your thoughts visual?” (1=completely nonvisual, 9=completely visual)
12. Auditory: “Were your thoughts verbal?” (1=completely nonverbal, 9=completely verbal)
13. Confidence: “How confident are you about your ratings for this trial?” (1=completely unconfident, 9=completely confident)

### EEG Preprocessing

We share both raw EEG data and preprocessed EEG data as derivatives. Preprocessing for all EEG data was run in MATLABR2024b^53^ using EEGLABv2024.2.1^54^ and Fieldtrip^55^ toolboxes. Preprocessing for each run was performed using the following pipeline: (1) In EEGLAB, the continuous EEG data were down sampled from 5000Hz to 250Hz; (2) Correction for MR gradient induced artifacts was performed using the BERGEN toolbox^56^ in EEGLAB. This toolbox relies on fMRI realignment parameters (RPs) obtained through motion correction, to calculate EEG artifact templates which are then subtracted from the raw data. To obtain the RPs, fMRI motion correction was performed with Statistical Parametric Mapping software (SPM12) in MATLAB^57^. For each run, ‘spm_realign’ was used to realign the fMRI volumes and obtain the six head-motion parameters (three translations and three rotations) for each volume. Motion parameters were thresholded at 0.8mm so that head movement exceeding 0.8mm was not included in the realignment template. The final two TRs were removed from the BOLD RP and EEG data because the EEG recording did not always contain sufficient data after the final fMRI volumes. Additional adjustments were made when the number of EEG triggers and fMRI volumes did not initially match. After synchronization, the fMRI motion parameters were passed to the Bergen MR correction toolbox to generate a motion-informed weighting matrix. The matrix, TR event markers, and a 500 sample (2-second) artifact window, were used as inputs to perform baseline correction and remove the MRI gradient artifact from the EEG. (3) Following gradient correction, the EEG data were corrected for cardioballistic and helium pump artifacts using ‘pop_cwregression,’ the CWL regression plugin for EEGLAB^38^. The four carbon wire loop channels (33–36) were used as reference signals to correct EEG channels (1–31). The CWL signals were used to construct time-varying, windowed regressors using a Hanning taper.

Beyond removal of scanner artifacts through the steps described above, no additional preprocessing was done on the shared EEG data to allow for maximum flexibility in subsequent preprocessing and analysis that researchers may choose to pursue. The **Technical Validation** section, however, details additional preprocessing conducted for quality checking purposes only. These outputs were not included as derivatives in the dataset upload; therefore the shared data does not include the additional technical validation steps applied, including filtering, channel interpolation, or independent component analysis. Data were referenced to the reference channel, located at FCz. All event markers originally recorded during data collection are retained in the EEG data. Users may consider additional preprocessing steps, such as filtering, re-referencing, additional artifact rejection (e.g. independent component analysis), and/or channel interpolation.

## Data Records

### Data availability

The raw and processed EEG-fMRI data are deposited on OpenNeuro: https://doi.org/10.18112/openneuro.ds007216.v1.0.0.8 Group-level fMRI statistical maps of the GLMs run on the GradCPT data are deposited on NeuroVault: https://identifiers.org/neurovault.collection:24458.

### Data organization

Raw MRI data have been converted from DICOM to NIFTI using dcm2niix^59^. Imaging data were de-identified by removing facial features from T1w MRIs using the FSL deface tool with fractional intensity for BET set to 0.3 for conservative defacing^60^. The EEG data are in the BrainVision three-file format (.eeg,.vhdr, and.vmrk). All data, including fMRI, EEG, stimulus timing, behavioral task responses, and surveys are organized according to the Brain Imaging Data Structure (BIDS)^61^ format, providing standardized folder and file naming with a nested structure hierarchy. The parent folder, titled ‘A multi-session simultaneous EEG-fMRI dataset with online experience sampling’ contains all unprocessed data, derivatives, and descriptor files. Within the parent folder, the raw data are organized by subjects, then sessions. Each session folder contains (1) an ‘eeg’ folder for EEG recording files, event files, and electrode coordinate files; (2) an ‘fmap’ folder containing field map files and; (3) a ‘func’ folder containing fMRI and event files from the session. Additionally, the first session folder of each subject also contains an ‘anat’ folder with the subject’s defaced T1-w file. The 2D fast T1-weighted image collected during the second session was omitted from the dataset as its primary purpose was for registration during scanning and it is not recommended to use for formal preprocessing and analysis methods. Each data file is accompanied by a JSON metadata file.

All preprocessed EEG data are stored in a ‘derivatives’ folder, organized by subject and session. Within each session folder is an ‘eeg’ folder containing data corrected for MR gradient induced artifacts alone (files ending in ‘_Bergen’) and data corrected for both MR gradient induced and cardio ballistic and helium pump artifacts using CWL regression (files ending in ‘Bergen_CWreg’). Preprocessed EEG data are shared as.set and.fdt files, native to EEGLAB. Functional MRI derivatives are not included in the shared dataset given the heterogeneity of preprocessing approaches; we leave these decisions to the users. Statistical maps of the GLMs run on the GradCPT data as part of the **Technical Validation** analyses are provided in the ‘derivatives’ folder. We did not include derivatives from the remaining validation analyses in the shared dataset.

All task data for EEG and fMRI files have corresponding event files in TSV format containing stimulus timing and behavioral response data located in the ‘eeg’ and ‘func’ folders. Event files located in the ‘eeg’ folder describe timing for event markers (i.e., Sync Status, TR, and stimulus markers) appearing in the EEG data. Event files in the ‘func’ folder contain trial, timing, and response information from GradCPT and MDES tasks. Files associated with the MDES task have been corrected for recording errors described in the “**Usage Notes**” section below. All original rating files associated with the MDES task which have not been flagged for correction follow the BIDS-format naming convention (files ending in “events.tsv”). Where runs have been identified as consistent with a mind-blank episode according to our assessment, two TSV files are provided corresponding to the original, uncorrected rating files (ending in “events_og.tsv”) and the corrected files which are named following the BIDS-format (ending in “events.tsv”) and mark the first probe of the trial as “N/A”. A CSV file titled, “mb_att_allsubs.csv,” lists all trials in the dataset that could be affected by the recording error and is included in the parent folder. Survey data are scored and summarized in a TSV file within the parent folder. Missing survey responses are indicated in the TSV file with a score of-9.

Unthresholded group-level z-statistical maps of the GLMs run on the GradCPT data as part of the **Technical Validation** analyses are provided on NeuroVault, a repository for openly sharing statistical brain maps.^61^ Three statistical maps are provided in a collection titled “GradCPT GLM validation maps for a multi-session simultaneous EEG-fMRI dataset.” Each map is associated with a unique identifier. The map titled “Commission errors” represents BOLD activity associated with participants incorrect responses to trials of a mountain scene during the GradCPT (ID: 1081519). The map titled “Correct Omissions” represents BOLD activity associated with correct omissions (correctly withheld responses to mountain scenes) during the GradCPT (ID: 1081520). The map titled “Variance time course” represents BOLD activity associated with trial-by-trial variance during the GradCPT (ID: 1081521). We did not include derivatives from the remaining validation analyses in the shared dataset.

## Technical Validation

### EEG quality control

Quality metrics were based on a quality control pipeline developed by Delorme et al.^48^ to assess EEG data and are consistent with those adopted by Telesford et al.^62^ in assessing data collected during simultaneous EEG-fMRI. The quality control pipeline considers the percents of “good” channels, “good” data, and components identified from independent components analysis (ICA) classified as “brain-based” (**Figure 2c**). Following this pipeline, the EEGLAB plugin clean_rawdata (v2.0) was used to calculate number of unusable channels and percentage of good data in each run. Channel removal was based on more than five seconds of flatline activity, Pearson’s correlation of less than 0.8 with nearby channels, and line noise greater than four standard deviations relative to its signal. The clean_rawdata tool removes artifact contaminated data, like that introduced through body movement, via Artifact Subspace Reconstruction^63^, by identifying segments with variance greater than 20 times the variance of the calibration data. We ran ICA to identify artifacts via EEGLAB’s RunICA plugin. The ICLabel tool was used to identify components that were most likely to be sources of brain activity compared to all other potential signal sources (i.e. muscle, eye, heart, line noise, channel noise, or other). The number of brain-based components was compared to the total number of ICs identified to calculate the percentage of brain ICs. Quality metrics generated from our study were compared with reference to the quality of 30 EEG datasets (ranging from n=2 to n=384) collected outside of an MRI scanner, evaluated by Delorme et al.^48^ Overall, the percent of good channels fell within the same range observed in the comparison datasets with the majority of runs having 90-100% good channels. The percent of good data points (50-80%) in our study was slightly below that in the comparison datasets (60-90%). The percent of independent components attributed to brain activity (50-60%) was higher in our dataset than that in the comparison datasets (10-40%).

### fMRI quality control

Quality of functional MRI data was assessed by manually inspecting all images (no major artifacts were detected) and computing mean framewise displacement (FD) using the Motion Correction FMRIB’s Linear Image Registration Tool (MCFLIRT) in FMRIB Software Library (FSL)^59^ within each run of each task for each participant (**Figure 3**). The median FD value was 0.08mm (*SD* = 0.04mm), which is comparable to, or smaller than that in other MRI datasets of similar size^64^. Significant head movement flagged through visual inspection is noted in the dataset description file “EEG-fMRI_image03.csv” in the “comments_misc” column.

**Figure 3.**
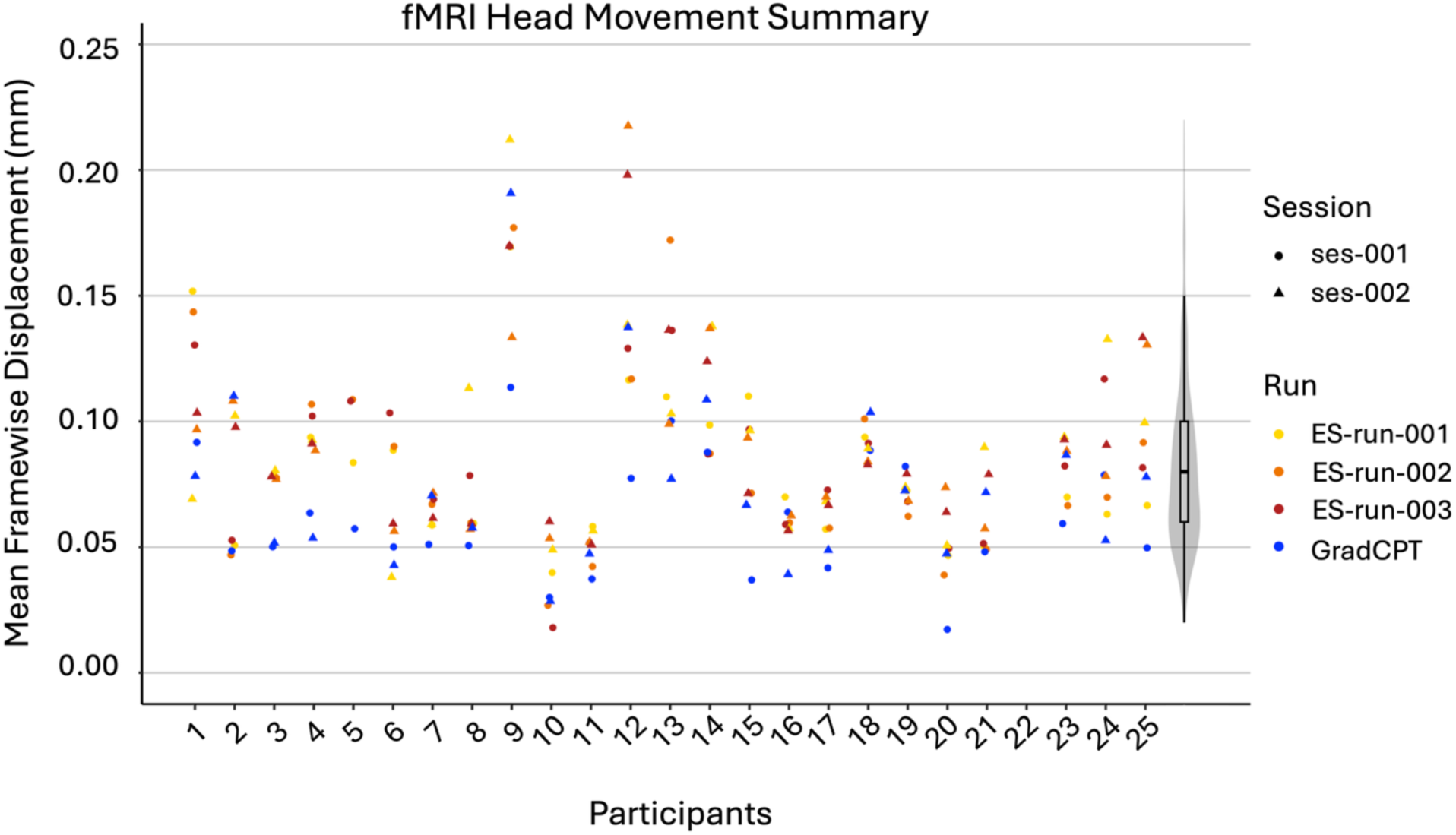
fMRI Head Movement Summary. Sessions are distinguished by data point shape (a circle indicating session 1 and a triangle indicating session 2), and task runs are distinguished by color (yellow, orange and red indicating the three runs of the rs-MDES task respectively and blue indicating GradCPT). Appended to the end of the graph is a violin plot in grey illustrating the overall distribution shape of the dataset. All mean FD values for all runs within the dataset fall below 0.25 mm.

To validate the suitability of the fMRI data for group-level brain–behavior analyses, we conducted whole-brain general linear model (GLM) analyses using FSL FEATv6.0^65,66^ with data collected from the GradCPT, given well-established prior fMRI findings from this task^32,67,68^. Due to data collection issues in one or both sessions of GradCPT, 4 subjects were excluded from analysis, leaving a total of 20 subjects (2 sessions each) suitable for GLM analysis. Prior to analysis, all fMRI runs were preprocessed in FEAT using MCFLIRT motion correction, spatial smoothing with a 5 mm full width at half maximum kernel, and a 100 s high-pass filter. Defaced T1s were skull-stripped using FSL’s brain extraction tool (BET) with a fractional intensity threshold set to 0.2 and biased field and neck cleanup. Boundary-based registration was used to align functional images to each subject’s respective T1. Linear normalization was then used to move from individual to standard MNI152 space.

Separate GLMs were performed to examine BOLD activation associated with correctly withheld responses to mountain scenes (correct omissions), commission errors, and trial-by-trial reaction time variance. To calculate a reaction time variance time course (VTC) within each run of GradCPT for each subject, we computed absolute *z*-scores of trial reaction time values. Similar to other recent studies, our approach takes all RTs into account, regardless of accuracy (omission trials were not included as they did not have associated reaction times).^65^ We normalized the scale so that each trial represented the deviation from the mean reaction time within the run. At the first level, all GLMs were computed for each run using double-gamma hemodynamic response function (HRF) with motion regressors included. Correct omission regressors were entered at the first level using event onset times for all trials where participants correctly withheld their response to a mountain scene. Commission error regressors were entered using event onset times for all trials where participants incorrectly responded to a trial of a mountain scene. VTC regressors were entered at the first level as trial-wise VTC values demeaned and HRF-convolved. Second-level models were run with fixed-effects within a single subject across all runs and sessions. A third-level model was computed for the full dataset using mixed-effects (FLAME 1). For each GLM, significant clusters were defined as those surviving a threshold of Z > 3.1 and cluster-level correction at p < 0.05. Network identification was based on the Yeo 7-network parcellation atlas in MNI152 2mm space^69^.

Our GLM analyses revealed significant positive and negative associations with correct omissions (**Figure 4a**) and commission errors (**Figure 4b)** at the group-level. Specifically, correct omissions were positively associated with nodes of the salience/ventral attention network (dorsal anterior insular cortex, supplementary motor area), somatomotor network (precentral), visual network (extrastriate cortex), and dorsal attention network (temporo-occipital cortex, superior parietal lobule). Nodes of the default network (inferior parietal lobule, posterior cingulate cortex) were negatively associated with correct omissions. Commission errors were positively associated with nodes of the salience/ventral attention network (dorsal anterior insular cortex, anterior midcingulate cortex) and were negatively associated with nodes of the default mode network (inferior parietal lobule, dorsomedial prefrontal cortex, lateral temporal cortex) and somatomotor network (post central). The network activation patterns we observed for both omission trials and commission errors are largely consistent with prior work^67^.

**Figure 4.**
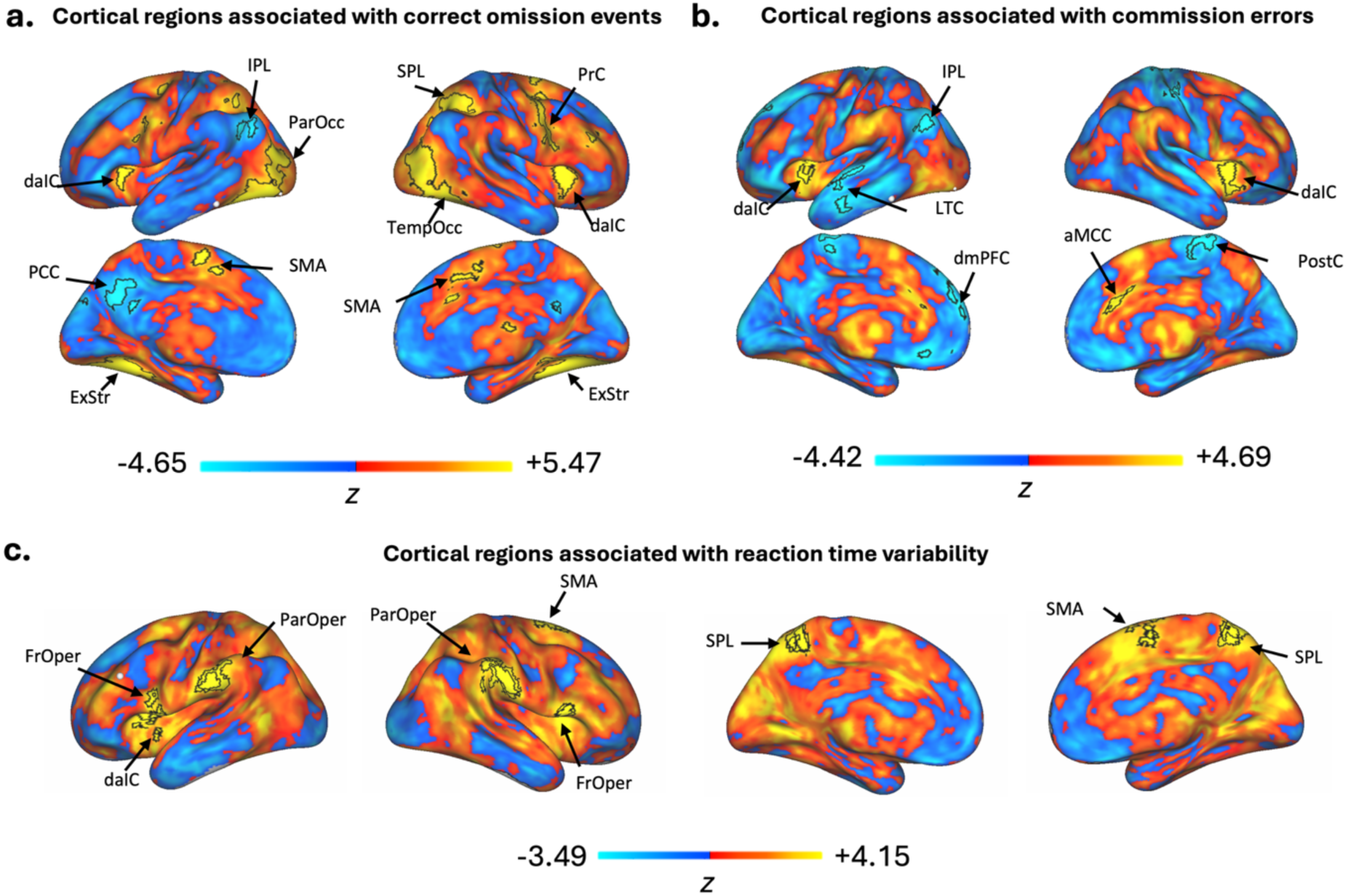
Group-level General Linear Model Results for GradCPT task. Unthresholded z-statistic maps showing positive (warm colors) and negative (cool colors) associations. Group-level results were visualized on the cortical surface using Connectome Workbench by projecting unthresholded z-statistic maps to the fsaverage6 surface. Black outlines indicate clusters surviving a cluster-forming threshold of Z > 3.1 and cluster-level correction at p < 0.05. Statistical maps were estimated using a mixed-effects general linear model (GLM). Maps are based on whole-brain, group-level (n=20) GLM analyses combining all GradCPT runs and sessions. (A) Regions activated during correct omissions. Yellow indicates areas positively associated with correct omissions. Blue indicates areas negatively associated with correct omissions. (B) Regions activated during commission errors. Yellow indicates areas positively associated with commission error events Blue indicates areas negatively associated with commission errors. (C) BOLD associations with the reaction time variance (RTV). Yellow color indicates areas positively associated with *higher reaction time variability during task, blue color indicates areas negatively associated with reaction time variability. Parietal Operculum (ParOper), supplementary motor area (SMA), Frontal operculum (FrOper), dorsal anterior insular cortex (daIC), anterior midcingulate cortex (aMCC), precentral (PrC), parietal occipital (ParOcc), temporal occipital (TempOcc), superior parietal lobule (SPL), post central (PostC), extrastriate cortex (ExStr), inferior parietal lobule (IPL), posterior cingulate cortex (PCC), inferior parietal lobule (IPL), dorsal medial prefrontal cortex (dmPFC), lateral temporal cortex (LTC)*

VTC analysis revealed significant positive BOLD associations, but not significant negative associations, at the group-level (**Figure 4c**). Networks showing significant correlations with VTC include the salience/ventral attention network (including the parietal operculum, frontal medial, frontal operculum, and insula) and dorsal attention network (superior parietal lobule). Positive associations with the salience and dorsal attention network shown in our data trend in the same direction as previously demonstrated GradCPT based BOLD associations^67,68^ with fluctuations in reaction time variance.

### Behavioral performance

The behavioral data from our tasks were analyzed using a variety of different quality metrics. Forty-three sessions contained GradCPT data, which were assessed for accuracy through commission and omission error rates to determine whether participants performed at an expected level (**Table 1**)^32^. The proportion of trials in which the participant incorrectly missed a response (i.e., to city stimuli) were counted as errors of omission (*M* = 0.05, *SD* = 0.05). Omission error rates within this dataset were within the typical range found in past studies (∼3-6%)^32,70^. The proportion of trials in which the participant incorrectly proceeded with a button press (i.e., to mountain stimuli) were counted as errors of commission (*M* = 0.38, *SD* = 0.18). Commission error rates within this dataset were somewhat higher than typical rates across past studies, which average around 25-30% in healthy adults^32,70^. This deviation from typical rates may be due to sampling variability and/or genuine behavioral factors We did not observe clear effects of time on accuracy, so an argument for decreased vigilance over time is not well supported. This heightened error rate is also not isolated to a few outlier subjects bringing the mean up but appears to be pervasive throughout the dataset. One possible explanation for these findings is that for the 8-minutes of the GradCPT task, participants remained in an “out of the zone” attentional state, where participants are effortfully exerting control over their attention to stay on task but is associated with higher levels of reaction time variability and commission errors.^32^ This hypothesis may be supported by the fact that the GradCPT task was administered as the first task in the study for each session, therefore participants may have been in a higher state of arousal which could lead to being “out of the zone.” Additionally, the instructions to respond “as quickly and accurately as possible” may have encouraged people to respond faster but with more mistakes. Another possibility could be high levels of distraction and mind-wandering in our participants. This cannot be verified, however, as we did not include thought-probes during the task to assess attention.

**Table 1.** Descriptive Data for the Gradual Onset Continuous Performance Task. All three metrics were calculated by averaging within each participant (across two runs), then across all participants. CE = Commission Error; OE = Omission Error; RT = Reaction Time (seconds).

|  | CE rate | OE rate | RT |
| --- | --- | --- | --- |
| Mean | 0.38 | 0.05 | 0.81 |
| SD | 0.18 | 0.05 | 0.05 |
| Range | 0.12 – 0.83 | 0.00 – 0.19 | 0.71-0.95 |

Resting state MDES ratings were assessed using group-level principal component analysis (PCA) with a 90-degree varimax rotation across all 13 thought probe dimensions to comprise a summary of how different thought dimensions group together and explain the variability across participants. PCA is a commonly used dimension reduction method in MDES studies to extract core thought types from the full set of items probed^6^. The Kaiser–Meyer–Olkin measure of sampling adequacy was 0.60, which surpasses the 0.50 threshold that is considered to be unacceptable, and Bartlett’s test of sphericity was significant [χ2(78) = 1460.47, *p* < 0.001], indicating the resulting dimensions provide an accurate description of the initial data. Twenty-five rs-MDES trials were excluded from analysis due to missing responses, leaving 815 trials which contributed to the PCA (96.91% of full dataset). We identified five components which explained 60% of the variability in ratings (**Figure 5)**. Several of these components show similar trends to past studies such as distinct components with paired positive loadings of “Others” and “Past” (PC1) and “Self” and “Future” (PC4)^71,72^, as well as an inner speech component (PC5) with high positive loading of the “Auditory” item and a high negative loading of the “Imagery” item^21^.

**Figure 5.**
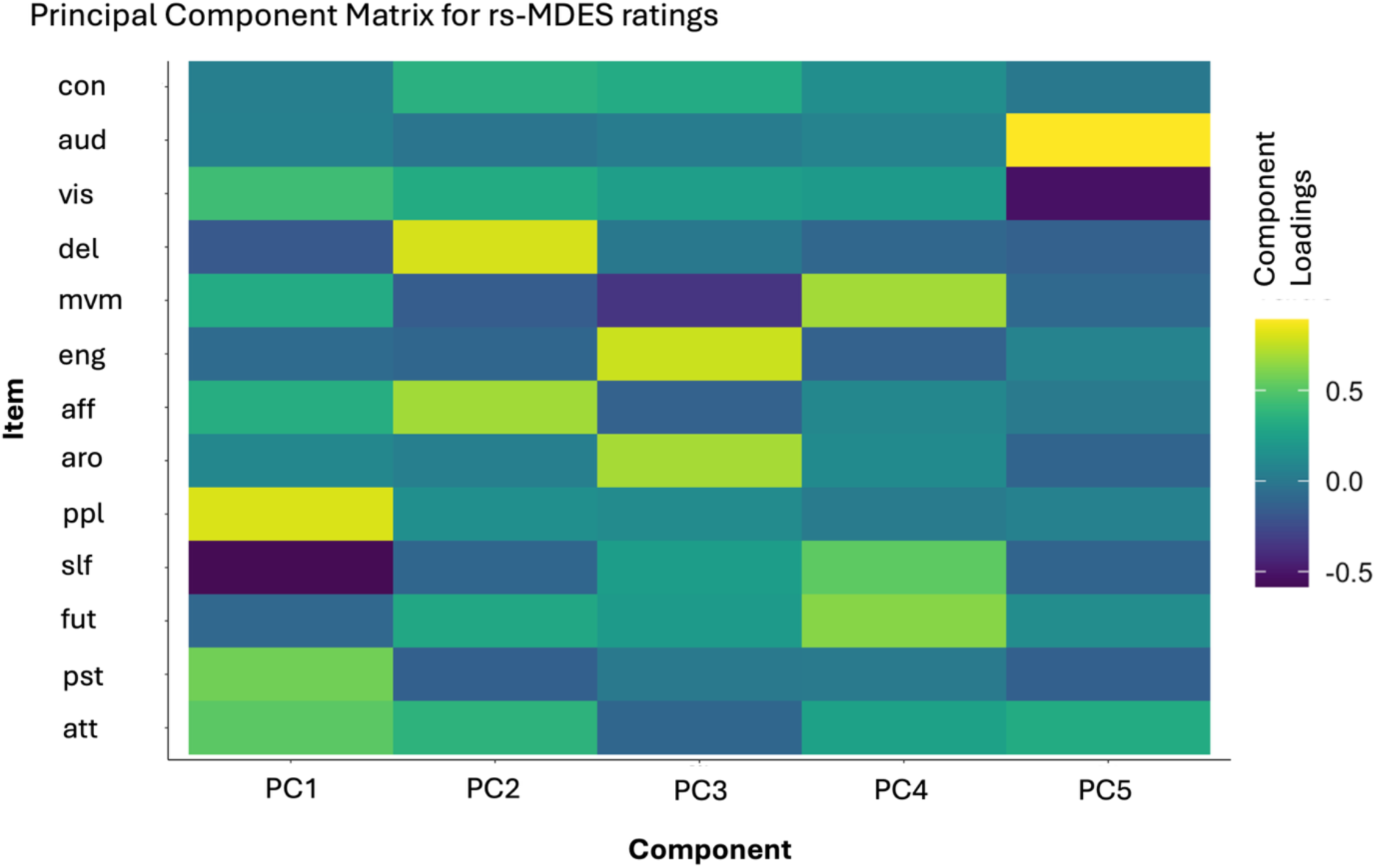
Principal component matrix for rs-MDES ratings. **a)** Heat map of principal components identified using all trials of data available and all 13 dimensions. Items: att = attention (where higher loadings correspond to more mental experiences); pst = past (where higher loadings correspond to more past oriented thinking); fut = future (where higher loadings correspond to more future oriented thinking); slf = self-relevant (where higher loadings correspond to more self-relevant thought); ppl = thoughts about other people (where higher loadings correspond to more thinking related to other people); aro = arousal (where higher loadings correspond to higher arousal); aff = affect (where higher loadings correspond to more positive affect); eng = engagement (where higher loadings correspond to higher engagement); mvm = movement (where higher loadings correspond to more freely moving thoughts); del = deliberate (where higher loadings correspond to more intentional thinking); vis = visual (where higher loadings correspond to more visual experiences); aud = auditory (where higher loadings correspond to more auditory experiences); con = confidence (where higher loadings correspond to higher confidence in rating).

Descriptives of the dimensions for the group were combined into Table 2 which includes the mean rating for each item, the standard deviation, and the number of trials included in each (excluding time-out trials and missing responses). Our rs-MDES paradigm also had a built-in quality check in our last item, which asks participants to rate their confidence in their responses to all other probes^30,73^. The high mean and small standard deviation for this item provides reassurance that most trials within the task can be considered “valid” as rated by the participants themselves (**Table 2**). In total, there were only 14 trials with confidence ratings below 5 in the whole dataset.

**Table 2.**
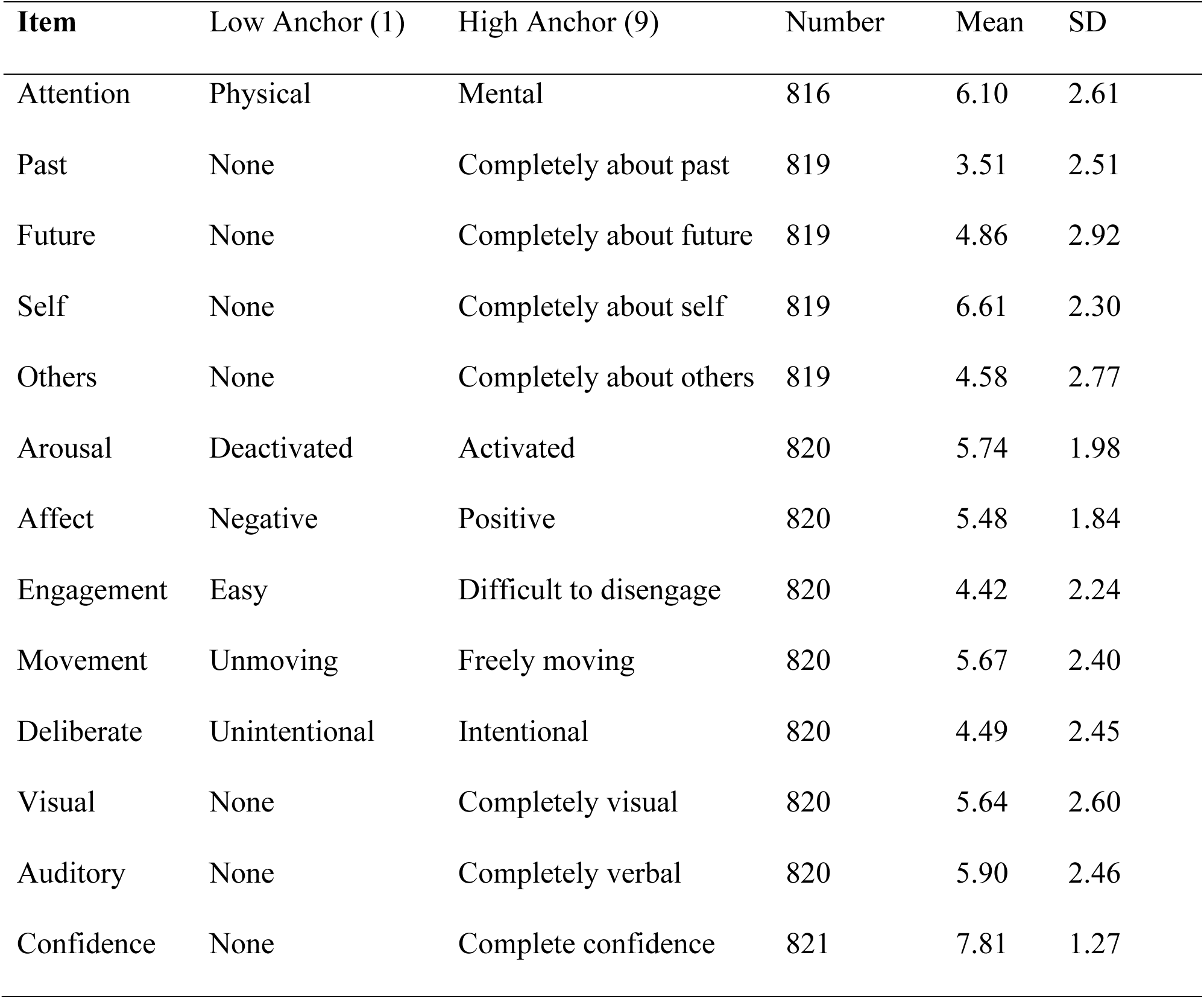
Descriptive Data for Repeated Multi-Dimensional Resting State Experience Sampling Ratings. For each item probed, the table describes the low and high anchors on the 1-9 Likert scale provided for participants, the number of complete ratings, and the mean and standard deviation for the item across all trials for all participants.

## Usage Notes

Researchers using the dataset should note that the 2D fast T1-weighted image collected during the second session was omitted from the dataset as its primary purpose was for field of view placement during scanning and it is not recommended to use for formal preprocessing and analysis methods. It is recommended that the 3D high resolution MPRAGE structural T1-weighted image from session one is used for preprocessing and analysis across all sessions collected within a single subject.

During data collection across the two sessions of 24 participants, various obstacles and malfunctions occurred. Included in the dataset description file (titled EEG-fMRI_image03.csv) we describe deviations from the standard procedure outline above in the “comments_misc” column. This will be necessary to inform any analysis performed on the data and give researchers a clearer and more transparent understanding of the data as they interact with it. The usage notes CSV file is available with the data and specifies which subject and session an issue pertains to (and describes the issue in detail).

One source of deviation stemmed from obstacles at data collection during the GradCPT task. Issues with the task ranged from failure to record participant responses (sub-001_ses-002; sub-025_ses-001) to extreme variation in the inter stimulus interval between trials due to stimulus presentation timing errors (sub-001_ses-001; sub-017_ses-002). This interfered with the data collection in a total of four GradCPT runs, leaving the final dataset with task response data from 43 out of 47 sessions.

Another issue arose in the rs-MDES task as there were bugs in the code for our stimulus which misrecorded participant ratings in two main ways. (1) The Likert scale for each question displayed only options from 1-9 which was the intended range for the task. It was possible, however, for a participant to advance the cursor off the scale to the extreme right (past 9) and enter a response that was recorded in the data as “10.” These responses were manually corrected to a rating of “9” which should be the most extreme response possible. (2) Mind-blank ratings for the first probe (“Stimulus-independent attention”) were erroneously recorded as a rating of 5 on the 1-9 scale between physical and mental thoughts (item #1). Our intent had been to record mind-‘blanking instances as a number that did not fall within the scale range in order to distinguish these instances from clear sensory and/or mental experiences. In total, 20 trials were determined to be either mind-blank experiences or were ambiguous (2.38% of all trials collected). To address the issue of conflated mind blanking ratings with ratings of 5 on the attention scale, we assessed each set of ratings that included an attention rating of 5 and created rules for exclusion based on what we determined to be most likely true mind-blank ratings, as follows: (1) a confidence rating less than 5 AND/OR (2) all content questions rated uniformly with a 5 or a 1. Justification for rule #1 was that if the confidence of a report is low enough, then the report is unreliable which may indicate that the scales did not meaningfully capture the experience, such as would be the case in an instance of mind blanking. Justification for rule #2 is that the rating of five on any scale is the default (where the cursor begins), therefore, if ratings of content specifically are all rated as 5, this may indicate that the participant quickly skipped through the questions after indicating a mind blank with the first probe, or, if all content is rated as 1 (i.e., not thinking at all about any content) this may be reflective of ratings congruent with the experience of mind blanking. If a trial was determined to be a mind-blank trial or was ambiguous, then the first “attention” rating value was changed to a NaN value. We include both corrected and original behavioral rating files on OpenNeuro.

Another important usage note concerns fMRI volume markers in the EEG data. The length of the rs-MDES task varied from run to run depending on the participant’s pace and the randomized duration of the fixation cross across trials. This necessitated manual termination of the scan upon conclusion of each rs-MDES run. In the majority of cases, the forced stop would end the functional scan in the middle of the collection of a volume, meaning that the EEG recording would receive an extra fMRI volume marker appended to the end of the recording; however, the scanner would not complete the measurement. This appended an extra fMRI marker (“T 1”) to the end of the EEG data which did not correspond to an fMRI volume. Exceptions to this rule are two runs which had two additional fMRI markers on the end of the EEG recording (sub-013_ses-002_run-003; sub-019_ses-001_run-003), four runs where there was one less fMRI marker in the EEG data than fMRI volumes recorded (sub-001_ses-002_run-003; sub-023_ses-002 all runs), and eight runs where the recorded fMRI markers and volumes were correctly aligned (sub-002_ses-001_run-002, run-003; sub-003_ses-002_run-001; sub-008_ses-001_run-001; sub-008_ses-002_run-002; sub-009_ses-002_run-002; sub-010_ses-001_run-001; sub-012_ses-002_run-003). In summary, this issue affects only the last markers in the run, where up to two volume markers in the EEG data may need to be omitted from analyses. All fMRI markers present in the EEG data are aligned with the first fMRI volume.

## Code Availability

The code for the resting state multi-dimensional experience sampling paradigm is available here: https://github.com/DynamicBrainMind/Experience-Sampling. The BIDS conversion tool used for this dataset is publicly available here: https://github.com/DynamicBrainMind/tobids. The EEG preprocessing pipeline used for this dataset is available here: https://github.com/DynamicBrainMind/eeg_fmri.

## Acknowledgements

All of the research reported in this publication was supported by the National Institute of Mental Health of the National Institutes of Health under award number R21MH127384 (to AK) and by the National Science Foundation Graduate Research Fellowship Program under Grant No DGE2439563 (to LST). The content is solely the responsibility of the authors and does not necessarily represent the official views of the National Institutes of Health or National Science Foundation.

## Author Contributions

Lotus Shareef-Trudeau contributed to study design, prepared the study protocol, collected data, performed technical validation analyses, and wrote the initial draft of the manuscript. David Braun contributed to dataset curation, technical validation analyses, and manuscript editing. Tiara Bounyarith assisted with data collection, data preparation, and technical validation analyses. Janet Li assisted with technical validation analyses and manuscript editing. Huiling Peng assisted with data collection, protocol planning and execution, and manuscript editing. Aaron Kucyi secured funding for the project, supervised and designed the project, assisted with protocol setup, assisted with technical validation analyses, and contributed to writing and editing of the manuscript.

## Competing Interests

The authors declare no competing interests.

## Notes

https://openneuro.org/datasets/ds007216

### Competing Interest Statement

The authors have declared no competing interest.

### Summary of Updates

Figure 2 was revised. The "Usage Notes" section was updated to clarify the contents of files shared for experience sampling ratings.

## References

1. Killingsworth, M. A. & Gilbert, D. T. A Wandering Mind Is an Unhappy Mind. Science 330, 932–932 (2010).

2. Seli, P. et al. How pervasive is mind wandering, really?,. Conscious. Cogn. 66, 74–78 (2018).

3. Kucyi, A., Kam, J. W. Y., Andrews-Hanna, J. R., Christoff, K. & Whitfield-Gabrieli, S. Recent advances in the neuroscience of spontaneous and off-task thought: implications for mental health. Nat. Ment. Health 1, 827–840 (2023).

4. Joshi, D., Tompary, A. & Kucyi, A. The default mode network: where spontaneous thought meets memory consolidation. Curr. Opin. Behav. Sci. 67, 101622 (2026).

5. Andrews-Hanna, J. R., Smallwood, J. & Spreng, R. N. The default network and self-generated thought: component processes, dynamic control, and clinical relevance. Ann. N. Y. Acad. Sci. 1316, 29–52 (2014).

6. Mills, C., Herrera-Bennett, A., Faber, M. & Christoff, K. Why the Mind Wanders. vol. 1 (Oxford University Press, 2018).

7. Smallwood, J. & Schooler, J. W. The Science of Mind Wandering: Empirically Navigating the Stream of Consciousness. Annu. Rev. Psychol. 66, 487–518 (2015).

8. Aaron Kucyi et al. A multi-session simultaneous EEG-fMRI dataset with online experience sampling. OpenNeuro 10.18112/OPENNEURO.DS007216.V1.0.0 (2026).

9. Groot, J. M. et al. Probing the neural signature of mind wandering with simultaneous fMRI-EEG and pupillometry. NeuroImage 224, 117412 (2021).

10. Chang, C. & Chen, J. E. Multimodal EEG-fMRI: Advancing insight into large-scale human brain dynamics. Curr. Opin. Biomed. Eng. 18, 100279 (2021).

11. Laufs, H. et al. Electroencephalographic signatures of attentional and cognitive default modes in spontaneous brain activity fluctuations at rest. Proc. Natl. Acad. Sci. 100, 11053–11058 (2003).

12. Mantini, D., Perrucci, M. G., Del Gratta, C., Romani, G. L. & Corbetta, M. Electrophysiological signatures of resting state networks in the human brain. Proc. Natl. Acad. Sci. 104, 13170–13175 (2007).

13. Xavier, M. et al. Consistency of resting-state correlations between fMRI networks and EEG band power. Imaging Neurosci. 3, IMAG.a.37 (2025).

14. Gonzalez-Castillo, J., Kam, J. W. Y., Hoy, C. W. & Bandettini, P. A. How to Interpret Resting-State fMRI: Ask Your Participants. J. Neurosci. 41, 1130–1141 (2021).

15. Kucyi, A. Just a thought: How mind-wandering is represented in dynamic brain connectivity. NeuroImage 180, 505–514 (2018).

16. Finn, E. S. Is it time to put rest to rest? Trends Cogn. Sci. 25, 1021–1032 (2021).

17. Lehmann, D., Henggeler, B., Koukkou, M. & Michel, C. M. Source localization of brain electric field frequency bands during conscious, spontaneous, visual imagery and abstract thought. *Cogn*. Brain Res. 1, 203–210 (1993).

18. Kam, J. W. Y. et al. Slow Fluctuations in Attentional Control of Sensory Cortex. J. Cogn. Neurosci. 23, 460–470 (2011).

19. Christoff, K., Gordon, A. M., Smallwood, J., Smith, R. & Schooler, J. W. Experience sampling during fMRI reveals default network and executive system contributions to mind wandering. Proc. Natl. Acad. Sci. 106, 8719–8724 (2009).

20. Martinon, L. M., Smallwood, J., McGann, D., Hamilton, C. & Riby, L. M. The disentanglement of the neural and experiential complexity of self-generated thoughts: A users guide to combining experience sampling with neuroimaging data. NeuroImage 192, 15–25 (2019).

21. Mckeown, B. et al. Self-reports map the landscape of task states derived from brain imaging. Commun. Psychol. 3, 8 (2025).

22. Kucyi, A. et al. Individual variability in neural representations of mind-wandering. Netw. Neurosci. 8, 808–836 (2024).

23. Van Calster, L., D’Argembeau, A., Salmon, E., Peters, F. & Majerus, S. Fluctuations of Attentional Networks and Default Mode Network during the Resting State Reflect Variations in Cognitive States: Evidence from a Novel Resting-state Experience Sampling Method. J. Cogn. Neurosci. 29, 95–113 (2017).

24. Vanhaudenhuyse, A. et al. Two Distinct Neuronal Networks Mediate the Awareness of Environment and of Self. J. Cogn. Neurosci. 23, 570–578 (2011).

25. Andrews-Hanna, J. R., Reidler, J. S., Huang, C. & Buckner, R. L. Evidence for the Default Network’s Role in Spontaneous Cognition. J. Neurophysiol. 104, 322–335 (2010).

26. Doucet, G. et al. Patterns of hemodynamic low-frequency oscillations in the brain are modulated by the nature of free thought during rest. NeuroImage 59, 3194–3200 (2012).

27. Mendes, N. et al. A functional connectome phenotyping dataset including cognitive state and personality measures. Sci. Data 6, 180307 (2019).

28. Braun, D. et al. Neural Sensitivity to the Heartbeat Is Modulated by Fluctuations in Affective Arousal during Spontaneous Thought. J. Neurosci. 45, e0608252025 (2025).

29. Kam, J. W. Y., Rahnuma, T., Jothiraj, S. N., Ouellette-Zuk, A. A. & Knight, R. T. Electrophysiological Signatures of Ongoing Thoughts during Naturalistic Behavior. Imaging Neurosci. (2025) doi:10.1162/IMAG.a.20.

30. Kane, M. J., Smeekens, B. A., Meier, M. E., Welhaf, M. S. & Phillips, N. E. Testing the construct validity of competing measurement approaches to probed mind-wandering reports. Behav. Res. Methods 53, 2372–2411 (2021).

31. Shareef-Trudeau, L., Lydon-Staley, D., Medaglia, J. & Kucyi, A. Brain-based predictive modeling of mind wandering and involuntary thought: The idiographic approach. Technol. Mind Behav. 6, (2025).

32. Esterman, M., Noonan, S. K., Rosenberg, M. & DeGutis, J. In the Zone or Zoning Out? Tracking Behavioral and Neural Fluctuations During Sustained Attention. Cereb. Cortex 23, 2712–2723 (2013).

33. Ives, J. R., Warach, S., Schmitt, F., Edelman, R. R. & Schomer, D. L. Monitoring the patient’s EEG during echo planar MRI. Electroencephalogr. Clin. Neurophysiol. 87, 417–420 (1993).

34. Laufs, H., Daunizeau, J., Carmichael, D. W. & Kleinschmidt, A. Recent advances in recording electrophysiological data simultaneously with magnetic resonance imaging. NeuroImage 40, 515–528 (2008).

35. Warbrick, T. Simultaneous EEG-fMRI: What Have We Learned and What Does the Future Hold? Sensors 22, 2262 (2022).

36. Luo, Q., Huang, X. & Glover, G. H. Ballistocardiogram artifact removal with a reference layer and standard EEG cap. J. Neurosci. Methods 233, 137–149 (2014).

37. Masterton, R. A. J., Abbott, D. F., Fleming, S. W. & Jackson, G. D. Measurement and reduction of motion and ballistocardiogram artefacts from simultaneous EEG and fMRI recordings. NeuroImage 37, 202–211 (2007).

38. Van Der Meer, J. N. et al. Carbon-wire loop based artifact correction outperforms post-processing EEG/fMRI corrections—A validation of a real-time simultaneous EEG/fMRI correction method. NeuroImage 125, 880–894 (2016).

39. American Psychiatric Association. DSM-5-TR Self-Rated Level 1 Cross-Cutting Symptom Measure—Adult.

40. Measuring Health and Disability: Manual for WHO Disability Assessment Schedule WHODAS 2.0. (World Health Organization, Geneva, 2010).

41. Spitzer, R. L., Kroenke, K., Williams, J. B. W. & Löwe, B. A Brief Measure for Assessing Generalized Anxiety Disorder: The GAD-7. Arch. Intern. Med. 166, 1092 (2006).

42. Kroenke, K., Spitzer, R. L. & Williams, J. B. W. The PHQ-9: Validity of a brief depression severity measure. J. Gen. Intern. Med. 16, 606–613 (2001).

43. Treynor, W., Gonzalez, R. & Nolen-Hoeksema, S. Rumination Reconsidered: A Psychometric Analysis. Cogn. Ther. Res. 27, 247–259 (2003).

44. Carriere, J. S. A., Seli, P. & Smilek, D. Wandering in both mind and body: Individual differences in mind wandering and inattention predict fidgeting. Can. J. Exp. Psychol. Rev. Can. Psychol. Expérimentale 67, 19–31 (2013).

45. Harris, P. A. et al. Research electronic data capture (REDCap)—A metadata-driven methodology and workflow process for providing translational research informatics support. J. Biomed. Inform. 42, 377–381 (2009).

46. Harris, P. A. et al. The REDCap consortium: Building an international community of software platform partners. J. Biomed. Inform. 95, 103208 (2019).

47. Speilberger, C. D., Gorsuch, Rl., Lushene, R., Vagg, P. & Jacobs, G. Manual for the state-trait anxiety inventory. Palo Alto CA Consult. Psychol. (1983).

48. Delorme, A. et al. Tools for Importing and Evaluating BIDS-EEG Formatted Data. in 2021 *10th International IEEE/EMBS Conference on Neural Engineering (NER)* 210–213 (IEEE, Italy, 2021). doi:10.1109/ner49283.2021.9441399.

49. Brainard, D. H. The Psychophysics Toolbox. Spat. Vis. 10, 433–436 (1997).

50. Hung, S.-M. & Hsieh, P.-J. Mind wandering in sensory cortices. Neuroimage Rep. 2, 100073 (2022).

51. Kane, M. J. et al. For Whom the Mind Wanders, and When, Varies Across Laboratory and Daily-Life Settings. Psychol. Sci. 28, 1271–1289 (2017).

52. Kucyi, A. et al. Prediction of stimulus-independent and task-unrelated thought from functional brain networks. Nat. Commun. 12, 1793 (2021).

53. The MathWorks Inc. MATLAB. The MathWorks Inc. (2024).

54. Delorme, A. & Makeig, S. EEGLAB: an open source toolbox for analysis of single-trial EEG dynamics including independent component analysis. J. Neurosci. Methods 134, 9–21 (2004).

55. Oostenveld, R., Fries, P., Maris, E. & Schoffelen, J.-M. FieldTrip: Open Source Software for Advanced Analysis of MEG, EEG, and Invasive Electrophysiological Data. Comput. Intell. Neurosci. 2011, 1–9 (2011).

56. Moosmann, M. et al. Realignment parameter-informed artefact correction for simultaneous EEG–fMRI recordings. NeuroImage 45, 1144–1150 (2009).

57. Statistical Parametric Mapping: The Analysis of Funtional Brain Images. (Elsevier/Academic Press, Amsterdam; Boston, 2007).

58. Li, X., Morgan, P. S., Ashburner, J., Smith, J. & Rorden, C. The first step for neuroimaging data analysis: DICOM to NIfTI conversion. J. Neurosci. Methods 264, 47–56 (2016).

59. Jenkinson, M., Beckmann, C. F., Behrens, T. E. J., Woolrich, M. W. & Smith, S. M. FSL. NeuroImage 62, 782–790 (2012).

60. Gorgolewski, K. J. et al. The brain imaging data structure, a format for organizing and describing outputs of neuroimaging experiments. Sci. Data 3, 160044 (2016).

61. Gorgolewski, K. J. et al. NeuroVault.org: a web-based repository for collecting and sharing unthresholded statistical maps of the human brain. *Front*. Neuroinformatics 9, (2015).

62. Telesford, Q. K. et al. An open-access dataset of naturalistic viewing using simultaneous EEG-fMRI. Sci. Data 10, 554 (2023).

63. Kothe, C. A. & Makeig, S. BCILAB: a platform for brain–computer interface development. J. Neural Eng. 10, 056014 (2013).

64. Jones, M. S., Zhu, Z., Bajracharya, A., Luor, A. & Peelle, J. E. A Multi-Dataset Evaluation of Frame Censoring for Motion Correction in Task-Based fMRI. Aperture Neuro 1–25 (2023) doi:10.52294/ApertureNeuro.2022.2.NXOR2026.

65. Woolrich, M. W., Ripley, B. D., Brady, M. & Smith, S. M. Temporal Autocorrelation in Univariate Linear Modeling of FMRI Data. NeuroImage 14, 1370–1386 (2001).

66. Woolrich, M. W., Behrens, T. E. J., Beckmann, C. F., Jenkinson, M. & Smith, S. M. Multilevel linear modelling for FMRI group analysis using Bayesian inference. NeuroImage 21, 1732–1747 (2004).

67. Fortenbaugh, F. C., Rothlein, D., McGlinchey, R., DeGutis, J. & Esterman, M. Tracking behavioral and neural fluctuations during sustained attention: A robust replication and extension. NeuroImage 171, 148–164 (2018).

68. Kondo, H. M. et al. Dynamic Transitions Between Brain States Predict Auditory Attentional Fluctuations. Front. Neurosci. 16, 816735 (2022).

69. Thomas Yeo, B. T., et al. The organization of the human cerebral cortex estimated by intrinsic functional connectivity. J. Neurophysiol. 106, 1125–1165 (2011).

70. Esterman, M., Rosenberg, M. D. & Noonan, S. K. Intrinsic Fluctuations in Sustained Attention and Distractor Processing. J. Neurosci. 34, 1724–1730 (2014).

71. Ruby, F. J. M., Smallwood, J., Engen, H. & Singer, T. How Self-Generated Thought Shapes Mood—The Relation between Mind-Wandering and Mood Depends on the Socio-Temporal Content of Thoughts. PLoS ONE 8, e77554 (2013).

72. Chitiz, L. et al. Mapping cognition across lab and daily life using Experience-Sampling. Conscious. Cogn. 131, 103853 (2025).

73. Polychroni, N., Herrojo Ruiz, M. & Terhune, D. B. Introspection confidence predicts EEG decoding of self-generated thoughts and meta-awareness. Hum. Brain Mapp. 43, 2311–2327 (2022).

